# Humanoid robots are perceived as an evolutionary threat

**DOI:** 10.1101/2021.08.13.456053

**Authors:** Zhengde Wei, Ying Chen, Jiecheng Ren, Piao Yi, Pengyu Zhang, Rujing Zha, Bensheng Qiu, Daren Zhang, Yanchao Bi, Shihui Han, Xiaochu Zhang

## Abstract

Studying how we perceive humanoid robots will provide insights for a deeper understanding of human-robot interaction. Our ideas about humanoid robots are mainly informed by science fiction, and humanoid robots are generally described as an evolutionary threat in science fiction that has not been tested. The preparedness model has emphasized that the fear module is automatically activated by evolutionary threats, and its underlying neural circuit is centered on the amygdala. We hypothesized that if humanoid robots are perceived as an evolutionary threat even though humanoid robots are manmade, modern objects, we would expect to observe a monocular advantage for humanoid robots and an amygdala response to unconsciously presented humanoid robots that were previously only evident in evolutionary threats. Here, we observed a monocular advantage for the perception of humanoid robots the same as an evolutionary threat (*i*.*e*., snakes). Our neuroimaging analysis indicated that unconscious presentation of humanoid robot vs. human images led to significant amygdala activation. Despite a positive humanoid robot-related association had been established by associative learning (as evidenced by results of successfully weakening the negative implicit attitude to humanoid robots and enhancing functional connectivity between the amygdala and hippocampus), the amygdala could still automatically and quickly detect humanoid robots. Our results reveal that processing of information about humanoid robots displays automaticity with regard to recruitment of visual pathway and amygdala activation. Our findings that humans apparently perceive humanoid robots as an evolutionary threat may help inform redefinition of human-robot interaction and robot ethics.

## Introduction

Artificial intelligence advances have led to robots that look and behave like humans. Studying how we perceive humanoid robots will provide insights for a deeper understanding of human-robot interaction, which may facilitate successful social encounters between humans and humanoid robots. A growing number of studies has found that the perceptions toward humanoid robots are complex and inconsistent (*1-5*). We contend that acquiring a better understanding of human perceptions toward humanoid robots would be facilitated by identifying what type of threat(s) humanoid robots are perceived as; insights in this area could help in redefining human-humanoid robot interaction and humanoid robot ethics.

From an evolutionary perspective, threats can be divided into evolutionary threats and modern threats (*6*). Evolutionary threats relate to potential life-threatening stimuli and situations frequently encountered in the environments of our early evolutionary ancestors. Modern threats are fear-relevant stimuli that poses a problem today that was not prevalent throughout human evolutionary history (*7*). Throughout human evolution, the ability to identify threatening stimuli has been critical to survival. Fear activates defensive behavior systems that help organisms to deal with different types of survival threats (*8, 9*). It has been proposed that fear is more likely to result from threat stimuli related to survival in evolutionary history. The preparedness model (*6, 10*) emphasizes that, compared to modern threats (*e*.*g*., weapons), processing of evolutionary threats (*e*.*g*., snakes) is assumed to show automaticity with regard to rapid recruitment of the behavioral and neural systems (see some cartoon demonstrations in Fig. 1). This automaticity persists despite the fact that modern threats may be equally or even more closely related with trauma in our daily lives. A monocular advantage in responses to a particular category may reflect visual facilitation of automatic processing for that category (*11*), and there is empirical evidence supporting that a monocular advantage is present in responses to images of evolutionary threats (*e*.*g*., snakes) but not in responses to images of modern threats (*e*.*g*., guns) (*11, 12*).

**Figure 1.**
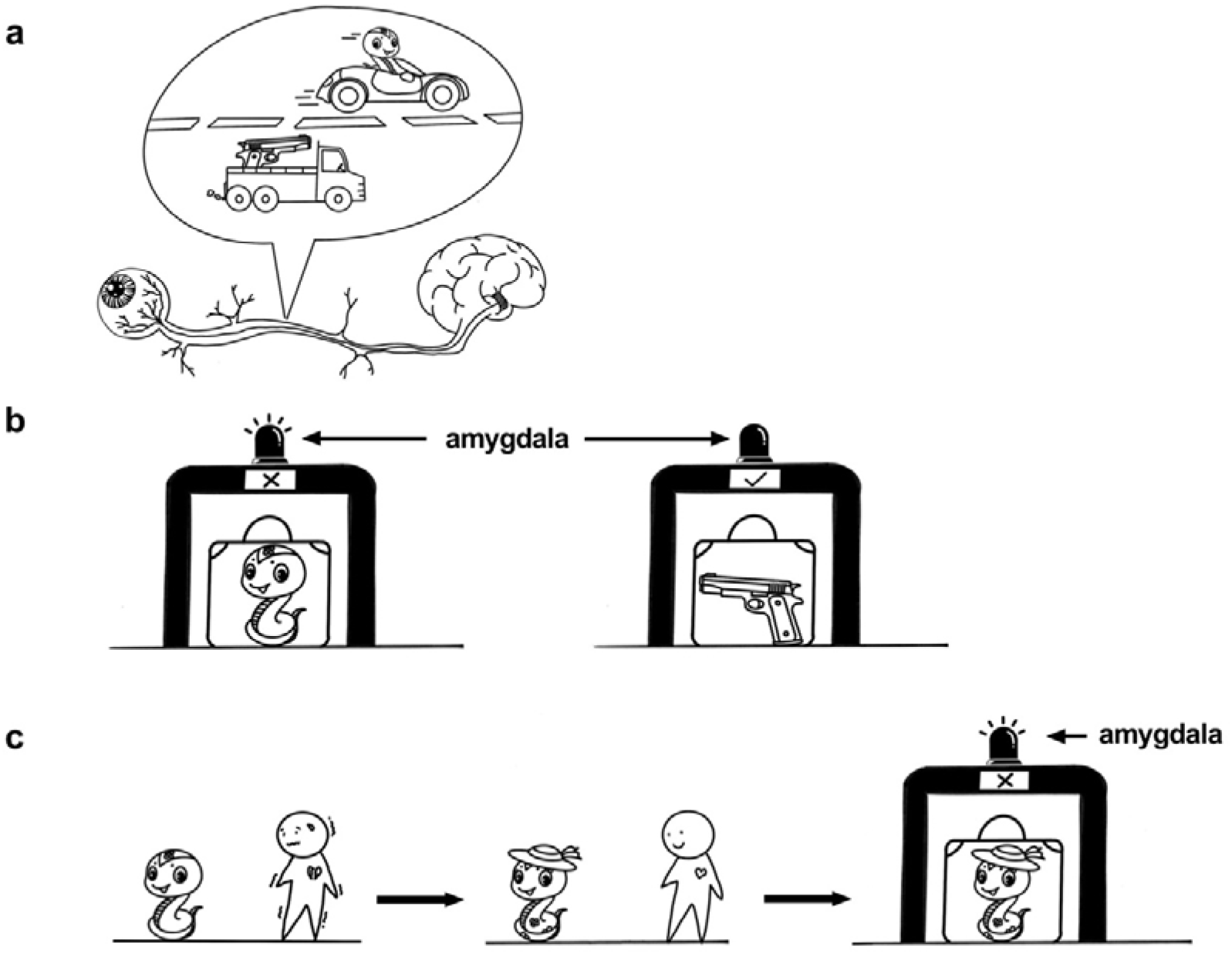
Automaticity for evolutionary threats as compared to modern threats. (a) Monocular advantage. The monocular input information of evolutionary threats (*e*.*g*., snake) is faster than information of modern threats (*e*.*g*., gun) to the brain for perception. (b) As the security devices can detect invisible dangerous goods in suitcases, the amygdala responds to unconsciously presented images of evolutionary threats, but do not respond to unconsciously presented images of modern threats. (c) Although we can turn our fear for evolutionary threats into explicit liking by some way (such as associative learning), the amygdala can still respond to unconsciously presented images of evolutionary threats.

The preparedness model (*6, 10*) also hypothesizes that the neural basis of automatic threat processing would be the amygdala. The amygdala has long been known to play a key role in responding to emotionally relevant stimuli, activating in response to images containing threatening or highly arousing features (*13, 14*). When automatic and controlled evaluations of threat sometimes differ, the more positive controlled processing can moderate more negative automatic processing (*15*), which could account for the absence of significant activation in amygdala in response to conscious presentations of threats (*16, 17*). Notable, although controlled processing can eliminate amygdala activation caused by consciously presented threats, amygdala still showed significant activation by threats when threats were presented unconsciously (*15-17*). The empirical evidence has demonstrated that amygdala responds more strongly to evolutionary threats than modern threats under unconscious presentation (*18*). It has also been proposed that there is enhanced resistance to extinction from evolutionary threats relative to modern threats (*10, 19*). Using classical conditioning paradigms, comparisons of aversive conditioning have been indicated enhanced resistance to extinction to snakes relative to guns (*19, 20*).

The humanoid robots are manmade, modern objects that have emerged recently in our cultural history. In previous studies, neither a monocular advantage (*11, 12*) nor an amygdala response (*21*) were detected upon encountering modern threats. Thus, we would anticipate that humanoid robots should fail to elicit a monocular advantage or an amygdala response. However, our ideas about humanoid robots are mainly informed by science fiction media (*22*). Science fiction has created the backdrop for how humanoid robots are interpreted and assessed (*22*). Humanoid robots are popular in science fiction (*23*) and are generally presented as artificial “living” entities with superhuman intelligence, digital emotions, and even consciousness(*22*). In many science fiction plots, these super-intelligent humanoid robots can exceed the abilities of human beings, and recent statements from some influential industry leaders have strengthened these fears (*24*). With the ideology that the intellectually superior are by nature masters and the intellectually inferior by nature slaves, it is comprehensible that we fear humanoid robots with superhuman intelligence will enslave us (*25*). So it is clear that some humanoid robots are perceived as more than “just tools”; rather, they are sometimes viewed as dangerous “living” entities. Thus, we proposed that humanoid robots are perceived as an evolutionary threat.

If humanoid robots are perceived as an evolutionary threat, then participants would be expected to show a monocular advantage for humanoid robot images and an amygdala response to unconscious presentation of humanoid robot images. Here, we pursued these topics by first measuring participants’ explicit and implicit attitudes to humanoid robots which revealed that despite positive attitudes found from self reporting in the questionnaire, participants actually had negative implicit attitudes to humanoid robots. We then used a Monocular Advantage Task which revealed that participants showed a monocular advantage for humanoid robots. We also employed fMRI and found that unconsciously presented images of humanoid robots trigger amygdala responses, and later there was resistance to changes in amygdala activity after successfully weakening the negative implicit attitudes of participants to humanoid robots. Our findings support the idea that humans may perceive humanoid robots as an evolutionary threat.

## Results

### Negative implicit attitudes to humanoid robots

People’s attitudes to humanoid robots are complex. In this study, we used an explicit attitude questionnaire (see the details in Supplementary Information) and the Implicit Association Test (IAT, Supplementary Fig. 1) to assess participants’ attitudes to humanoid robots. The IAT task reflects the degree of automatic connection between concept categories (*e*.*g*. humanoid or human) and attribute categories (*e*.*g*. threatening or non-threatening) by comparing the response time in combinations of “humanoid + threatening” and “human + non-threatening” and combinations of “humanoid + non-threatening” and “human + threatening”. If there is a stronger association between humanoid robots and threatening meaning than between humans and threatening meaning, then we would expect participants to respond faster when humanoid images and threatening words share the same response.

In the explicit attitude questionnaire, participants (n = 66, age: 21.65±2.38 years; 41 females) showed a positive explicit attitude toward humanoid robots (t_65_ = 8.84, p < 0.001, Cohen’s d = 1.10). In the IAT, participants (n = 127, age: 21.35±2.41 years; 80 females) had significant IAT scores (mean = 0.48; SD = 0.49; t_126_ = 10.95, p < 0.0001, Cohen’s d = 0.97; Fig. 2a). Participants responded faster in combinations of “humanoid + threatening” and “human + non-threatening” than in combinations of “humanoid + non-threatening” and “human + threatening” (t_126_ = -8.81, p < 0.0001, Cohen’s d = 0.78; Fig. 2a). Our IAT results indicated that, despite the findings from self reporting in the questionnaire, participants actually have negative implicit attitudes to humanoid robots.

**Figure 2.**
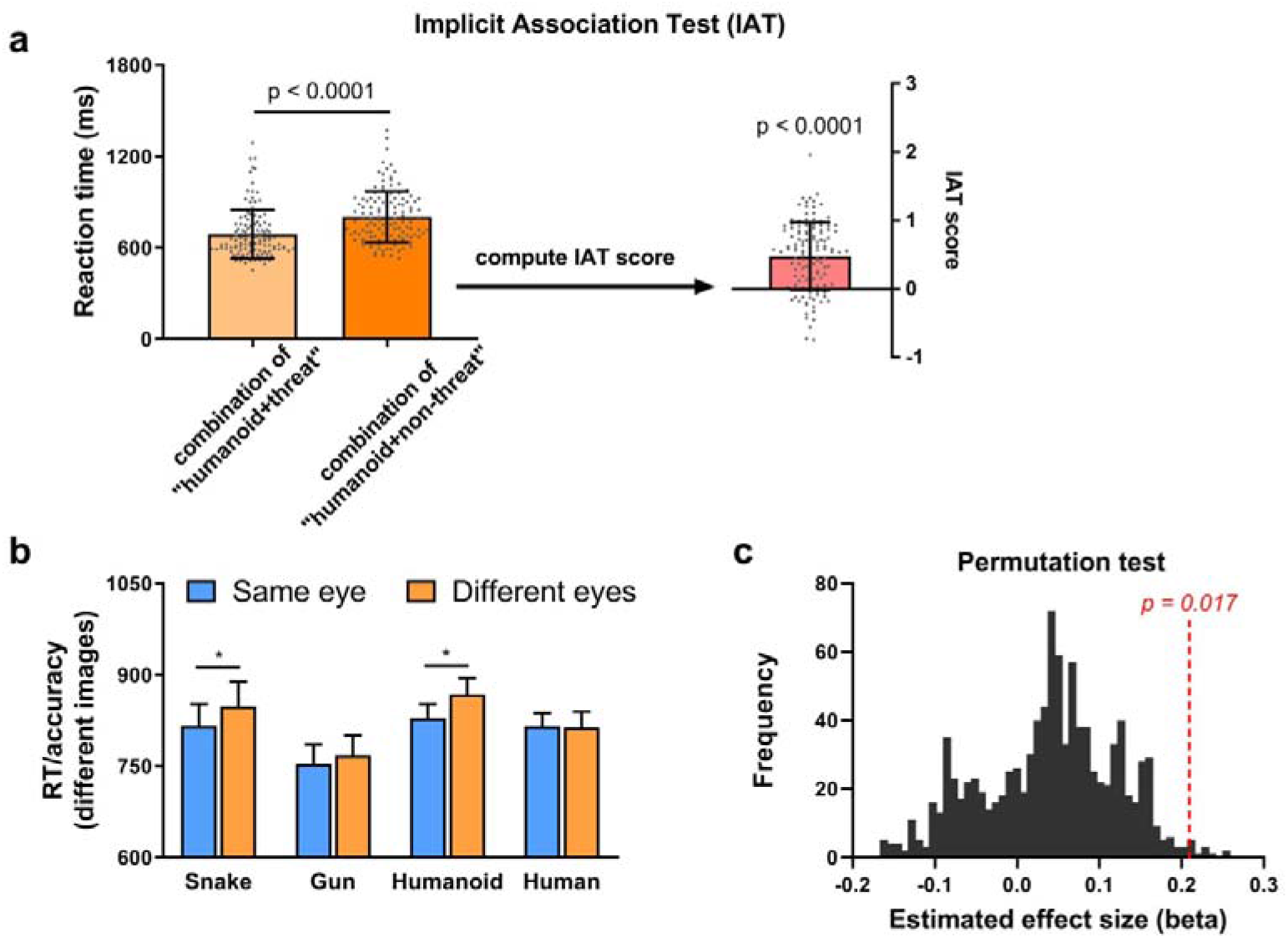
Implicit Association Test and Monocular Advantage Task. (a) Negative implicit attitude to humanoid robots. In the Implicit Association Test, participants responded faster in combination of “humanoid + threat” than in combination of “humanoid + non-threat”, and the computed IAT scores (effect size) were significant. Plotted data represent the mean ± SD across participants. (b) A monocular advantage for the perception of humanoid robot images and snake images. In the different condition of Image Match, the inverse efficiency (reaction time (RT)/accuracy) in the same eye input condition was smaller than in different eye input condition for humanoid robot images, and for snake images. Plotted data represent the mean ± s.e.m. across participants. (c) The inverse efficiency differences between Eye Input in the different condition of Image Match could successfully predict perceived threat category (perceived evolutionary threat (*i*.*e*., snake and humanoid robot) and perceived non-evolutionary threat (*i*.*e*., gun and human)). To test the significance of the effect size of prediction, a permutation test was performed. We exchanged the category labels of two randomly selected samples and calculated the effect size of the inverse efficiency differences, using a new general linear regression model each time. We repeated this step 1000 times and found that the original effect size survived the permutation test (p = 0.017). * p < 0.05.

Humanoid robots may be more intelligent and competitive than animal robots, resulting in more negative implicit attitudes to humanoid robots as compared to animal robots. We recruited another group of participants (n = 38, age: 21.77±1.67 years; 23 females) to perform a Humanoid-weapon IAT and a Animal robot-weapon IAT. Participants displayed larger IAT scores in Humanoid-weapon IAT than in Animal robot-weapon IAT (t_37_ = 3.07, p < 0.01, Cohen’s d = 0.50; Supplementary Figure 2; Supplementary Table 1). These results indicate that participants have a more negative implicit attitude to humanoid robots compared to animal robots.

### Monocular advantage for perception of humanoid robot images

A previous study indicated that a monocular advantage is shown in response to images of an evolutionary threat (*i*.*e*., snakes) but not in response to images of a modern threat (*i*.*e*., guns) (*11*). Here, we used a Monocular Advantage Task (Supplementary Figure 3) to investigate whether the visual pathway facilitates behavioral responses to humanoid robots. We hypothesized that if the humanoid robots are perceived as an evolutionary threat (as for snakes), then we would detect a monocular advantage for humanoid robots.

Thirty-two participants (age: 24.34±2.18 years; 21 females) were recruited. Each participant completed two blocks of trials for each of four categories (snake, gun, humanoid robot, and human), for a total of eight blocks. For each participant and condition, we measured mean accuracy and reaction time for each condition. We then computed inverse efficiency by dividing response time by accuracy (*11, 12*). We carried out a repeated-measures factorial ANOVA with Image Category (snake, gun, humanoid robot, and human), Image Match (same and different), and Eye Input (same and different eye) as within-subject factors, and with inverse efficiency as the dependent variable.

There was a significant main effect of Image Match (F(1,31) = 5.70, p = 0.017), no main effect of Image Category nor Eye Input (all p > 0.18), and no significant interaction (all p > 0.75). In the planned paired-sample t-tests, we compared the differences between same and different eye input for each image category. As shown in Fig. 2b, in the different condition of Image Match, the inverse efficiency in same condition of Eye Input was smaller than in different condition of Eye Input for snake images (t_31_ = -2.21, p = 0.035, Cohen’s d = 0.39), and for humanoid robot images (t_31_ = -2.23, p = 0.028, Cohen’s d = 0.41), but not for human images or gun images (all p > 0.24). In the same condition of Image Match, the inverse efficiency differences between Eye Input were not significant for any image category (all p > 0.10). These results indicate a monocular advantage for snakes and for humanoid robots.

We then used a general linear regression analysis to examine whether the inverse efficiency differences between Eye Input in the different condition of Image Match could successfully predict perceived threat category (perceived evolutionary threat (*i*.*e*., snake and humanoid robot) and perceived non-evolutionary threat (*i*.*e*., gun and human)). The inverse efficiency differences significantly predicted perceived threat category (t(126) = 2.185, p = 0.031, beta = 0.191, CI = [0.105, 2.122]). To further test the significance of the effect size (beta), a permutation test was performed. We exchanged the category labels of two randomly selected samples and calculated the effect size of the inverse efficiency differences, using a new general linear regression model each time. We repeated this step 1000 times and found that the original effect size survived the permutation test (p = 0.017, Fig. 2c).

### Greater amygdala activity induced by humanoid robot images compared to human images under unconscious presentation

Sixty-one participants (age: 21.03±2.43 years; 39 females) performed the IAT, and then they finished a modified Backward Masking Task (Fig. 3a-b) with fMRI scanning to measure amygdala activity in response to conscious and unconscious presentations of humanoid robot images. Following the end of fMRI scanning, all participants took part in a Forced-choice Detection Task to confirm that participants were aware of the stimulus under conscious presentation but were unaware under unconscious presentation in the Backward Masking Task. Twenty-five (age: 20.44±2.00 years; 16 females) out of the 61 participants performed the Evaluating Conditioning Task to weaken their negative implicit attitude to humanoid robots (the modulation effect was established in a pilot experiment, see the details in Supplementary Information, Supplementary Figure 4), and performed the modified Backward Masking Task with a second fMRI scanning. Following the end of fMRI scanning, participants took part in the IAT outside the scanner again (Fig. 3c).

**Figure 3.**
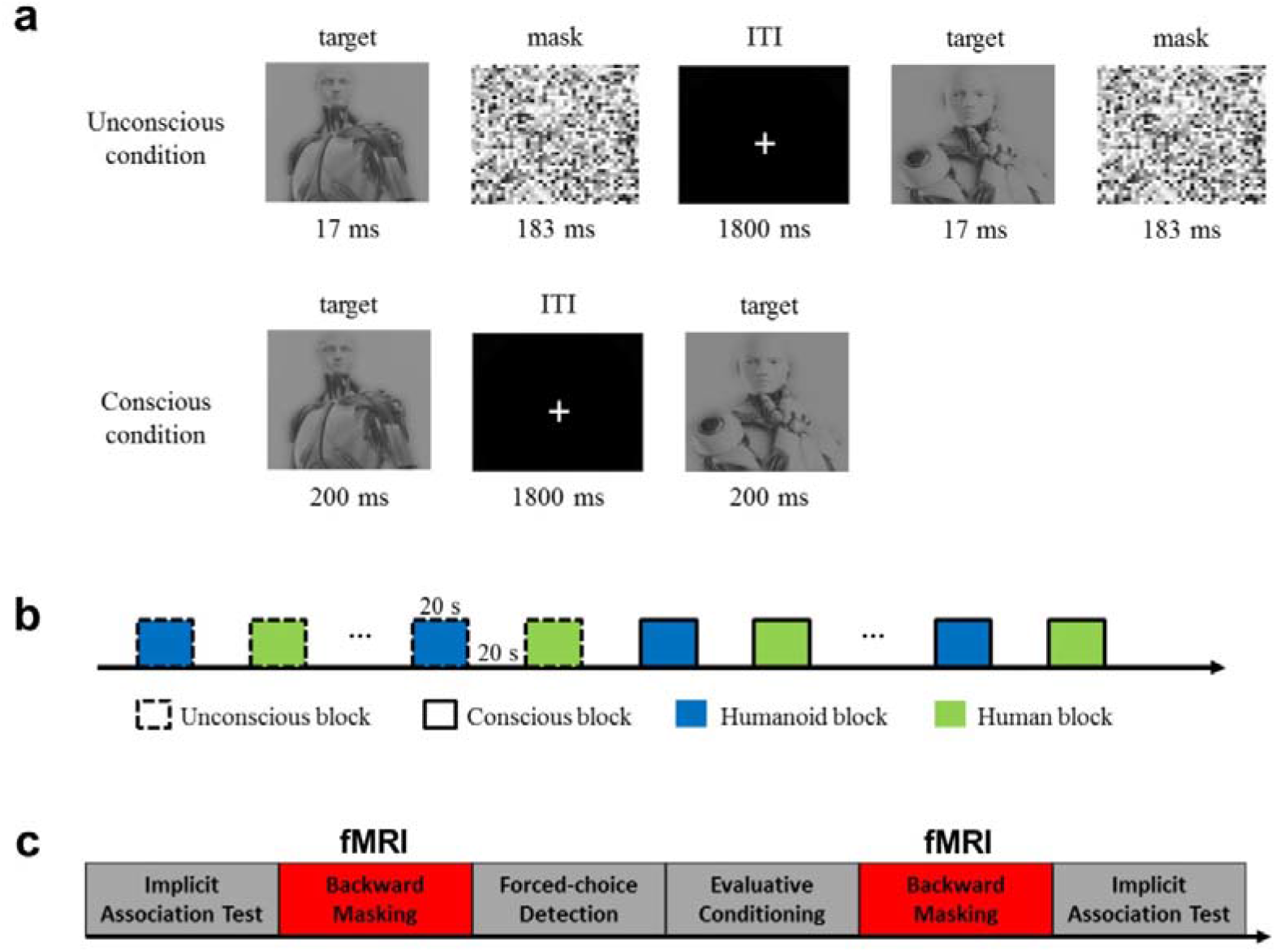
Description of the Backward Masking Task and the procedure for the fMRI experiment. (a) Time setting of the Backward Masking Task. In the unconscious condition, the target image was presented for 17 ms followed by a mask for 183 ms and a fixation for 1800 ms. In the conscious condition, the target image was presented for 200 ms followed by a fixation for 1800 ms. (b) Block design of the Backward Masking Task. There were six unconscious blocks (three humanoid blocks and three human blocks) followed by six conscious blocks (three humanoid blocks and three human blocks). (c) Procedure for the fMRI experiment. Participants performed an Implicit Association Test outside scanner, and then finished a modified Backward Mask Task with fMRI scanning followed by a Forced-choice Detection Task and an Evaluating Conditioning Task in scanner, and then they performed the modified Backward Mask Task with a second fMRI scan. Following the end of fMRI scanning, participants finished the Implicit Association Test outside scanner again.

As show in Supplementary Figure 5, the mean response rate was more than 90% under both presentations, and was higher under conscious presentation (t_58_= 4.14, p < 0.001, Cohen’s d = 0.54) than under unconscious presentation. The response time under conscious presentation was significantly shorter than that under unconscious presentation (t_58_= -10.91, p < 0.001, Cohen’s d = 1.42). The accuracy under conscious presentation was significantly higher than under unconscious presentation (t_58_= 16.68, p < 0.001, Cohen’s d = 2.17). Importantly, the accuracy under unconscious presentation did not differ from random chance (t_58_= - 0.31, p = 0.76), whereas the accuracy under conscious presentation was higher than random chance (t_58_= 19.13, p < 0.001, Cohen’s d = 2.49). These findings indicate that participants are aware of the stimuli under conscious presentation but are unaware of the stimuli under unconscious presentation.

In fMRI analysis, we first tested whether unconscious presentation of images of humanoid robots leads to greater amygdala response compared to images of humans. Activation in response to humanoid robot images was significantly stronger than in response to human images in anatomically defined bilateral amygdala (Fig. 4a) under unconscious presentation (left amygdala: t_60_ = 3.76, p < 0.001, Cohen’s d = 0.48; right amygdala: t_60_ = 3.32, p < 0.005, Cohen’s d = 0.42; Fig. 4b); no such difference was observed upon conscious presentation (left amygdala: t_60_ = -0.61, p = 0.55; right amygdala: t_60_ = -0.79, p = 0.43). The activation differences between humanoid robot and human images under unconscious presentation in bilateral amygdala were significantly stronger than activation differences under conscious presentation (left amygdala: t_60_ = 3.34, p < 0.001, Cohen’s d = 0.43; right amygdala: t_60_ = 2.67, p < 0.01, Cohen’s d = 0.34; Fig. 4b). Our results indicate that greater amygdala activity is induced by humanoid robot images compared to human images under unconscious presentation.

**Figure 4.**
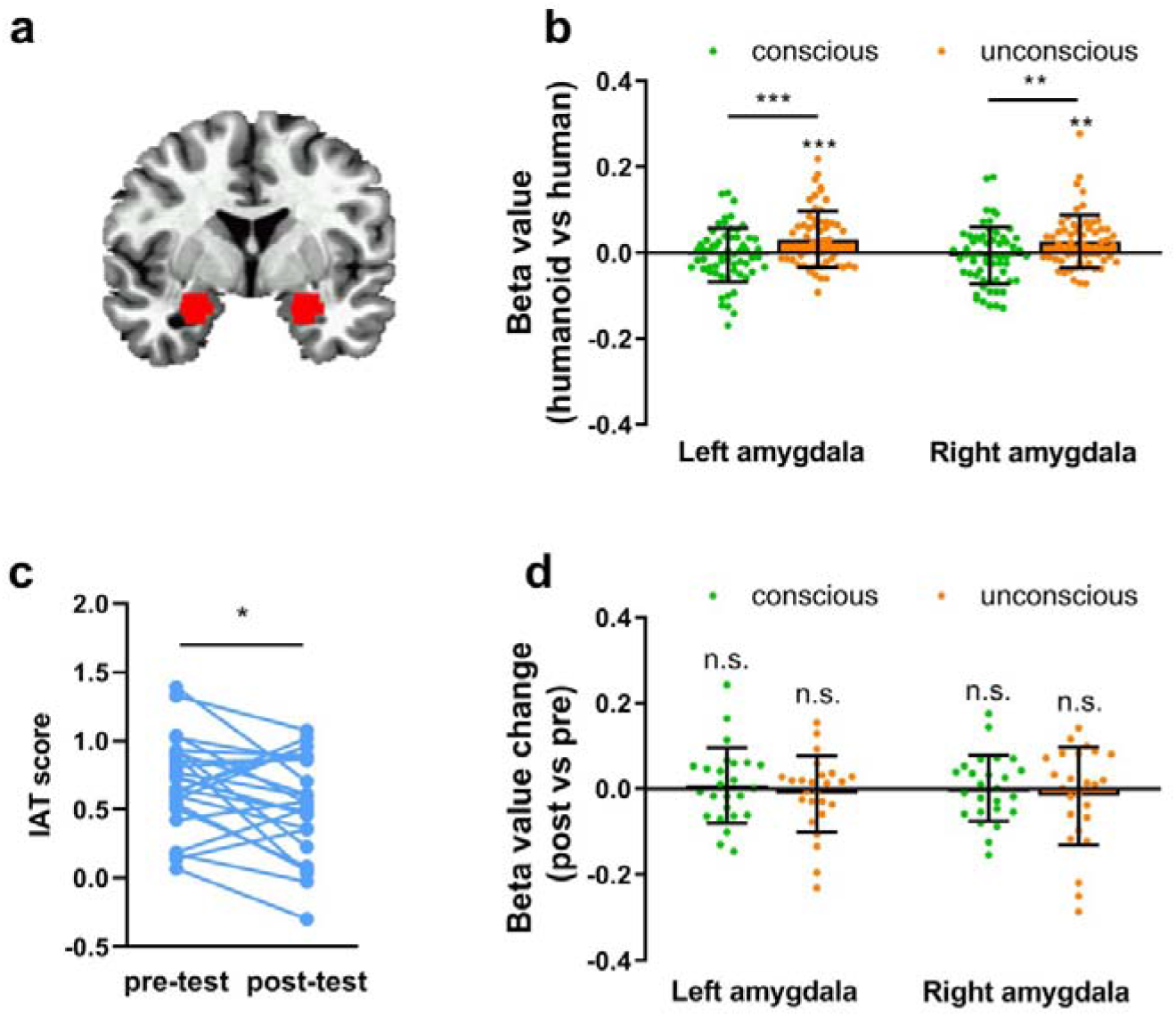
Humanoid robot image related amygdala activity and amygdala activity change. (a) The region of interest for bilateral amygdala. (b) Although no amygdala activity differences were detected for consciously presented humanoid robot vs. human images, greater amygdala activity was induced by humanoid robot images compared to images of humans under unconscious presentation. (c) Significantly smaller IAT scores were found in post-test compared to pre-test, indicating that participants’ negative implicit attitude to humanoid robots was successfully weakened. (d) Despite successfully weakening the negative implicit attitudes to humanoid robots, the amygdala activity differences of humanoid robot vs. human images did not change under conscious or unconscious presentations. * p < 0.05, ** p < 0.01, *** p < 0.001, n.s. = not significant. For a and c, plotted data represent the mean ± SD. across participants. IAT = implicit association test.

### Neural activity and functional connectivity changed after successfully weakening negative attitude

The Evaluative Conditioning Task was used to weaken the negative implicit attitude to humanoid robots. We found that the post-test IAT scores were significantly smaller than pre-test (t_24_ = - 2.46, p < 0.05, Cohen’s d = 0.49; Fig 4c), indicating that participants’ negative implicit attitude to humanoid robots had been successfully weakened by the Evaluative Conditioning Task. We then analyzed neuroimaging data for the change of neural activity and functional connectivity after successfully weakening negative attitude. A whole-brain analysis demonstrated a significant time main effect (pre-test, post-test) in the activation of the right DLPFC (uncorrected p < 0.005; Supplementary Figure 6). Previous studies defined ROIs of bilateral DLPFC as a sphere with a radius of 5mm centered at specific coordinates (Talairach coordinates; left: x = -47, y = 17, z = 28; right: x = 47, y = 20, z = 26) (*26*). Our ROI analysis revealed that bilateral DLPFC activation differences between humanoid robot and human images marginal significantly decreased in post-test compared to pre-test (left: t_24_ = 1.64, p = 0.06 (one-tailed), Cohen’s d = 0.33; right: t_24_ = 1.92, p < 0.05 (one-tailed), Cohen’s d = 0.38; Supplementary Figure 7a). Importantly, the activation change (post-test vs. pre-test) in the right DLPFC was positively correlated with IAT scores change (post-test vs. pre-test) (r = 0.40, p < 0.05; Supplementary Figure 7b).

We then used psychophysiological interaction analysis with a seed in the amygdala to test functional connectivity changes after successfully weakening negative attitudes. A whole-brain connectivity analysis demonstrated a significant time main effect in functional connectivity of the right amygdala and the right hippocampus (Fig. 5a), and thalamus (Supplementary Figure 8). We conducted a two factor (time factor: pre-test, post-test; image factor: humanoid robot, human) repeated measure ANOVA on functional connectivity under unconscious presentation of right amygdala-right hippocampus, and right amygdala-thalamus. There were significant time × image interaction effects in both couplings (amygdala-hippocampus: F_(1, 24)_ = 11.45, p < 0.001, Fig 5b; amygdala-thalamus: F_(1, 24)_ = 16.05, p < 0.001). There was a significant time main effect (F_(1, 24)_ = 10.47, p < 0.01) in the right amygdala-right hippocampus. There was a marginal significant image main effect (F_(1, 24)_ = 3.80, p = 0.054) in the right amygdala-thalamus. Importantly, the right amygdala-right hippocampus connectivity change (post-test vs. pre-test) was negatively correlated with the IAT scores change (post-test vs. pre-test) (r = -0.53, p < 0.01; Fig. 5c). These findings suggest that a positive humanoid robot-related association has been established by associative learning as evidenced by results of successfully weakening the negative implicit attitude to humanoid robots and enhancing functional connectivity between the amygdala and hippocampus.

**Figure 5.**
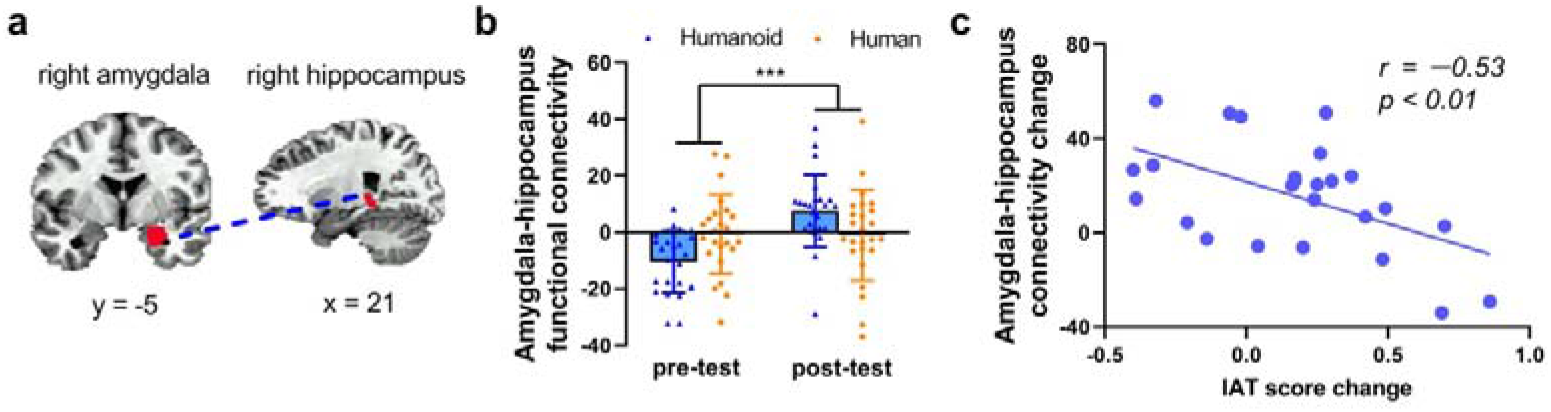
Amygdala-hippocampus functional connectivity was enhanced after attitude modulation. (a) A whole brain connectivity analysis demonstrated a significant time main effect in functional connectivity of right amygdala and right hippocampus. (b) Amygdala-hippocampus functional connectivity was enhanced in the post-test compared to the pre-test. We conducted a two factor (time factor: pre-test, post-test; image factor: humanoid, human) repeated measure ANOVA on amygdala-hippocampus functional connectivity under unconscious presentation. There was a significant time × image interaction effect (F_(1, 24)_ = 11.45, p < 0.001) and a significant time main effect (F_(1, 24)_ = 10.47, p < 0.01), but no significant image main effect (F_(1,24)_ = 0.51, p = 0.82). (c) IAT score change was negatively correlated with the change of amygdala-hippocampus functional connectivity (r = -0.53, p < 0.01). For b, plotted data represent the mean ± SD. across participants. IAT = Implicit Association Test.

### The amygdala response to unconscious presentation of humanoid robot images does not change after successfully weakening negative attitude

We next tested whether the amygdala response to unconscious presentation of humanoid robot images changes after successful weakening negative implicit attitude. We conducted a two factor (time factor: pre-test, post-test; presentation factor: unconscious presentation, conscious presentation) repeated measure ANOVA on activation differences between humanoid robot and human images in the left and right amygdala. No significant time×presentation interaction effects were found in any brain region (all p > 0.39). There were no significant time or presentation main effects in the left or right amygdala (all p > 0.55). *Post-hoc* analysis revealed no changes in bilateral amygdala activation between humanoid robot and human images under the conscious or unconscious presentations between the post-test and pre-test scans (all p > 0.48; Fig. 4d). Correlation analyses revealed that there was a significant correlation between IAT scores change and activation value change in the left amygdala under conscious presentation (r = 0.39, p = 0.050). However, we found no significant correlation between IAT scores change and activation value change in bilateral amygdala under unconscious presentation (all p > 0.13). These results demonstrate that despite a positive humanoid robot-related association had been established by associative learning, the amygdala could still automatically and quickly detect humanoid robots.

## Discussion

We found automaticity for processing information about humanoid robots that were previously only evident in evolutionary threats. The automaticity is supported by a monocular advantage for humanoid robots and an amygdala response to unconsciously presented humanoid robots. The monocular advantage in responses to humanoid robots reflects visual facilitation of responses for humanoid robots. The amygdala response to unconsciously presented humanoid robots indicates the automatic and quick detection of humanoid robots. Despite a positive humanoid robot-related association has been established by associative learning, the amygdala can still automatically and quickly detect humanoid robots.

Our results provided evidence for the idea that humanoid robots may be perceived as an evolutionary threat. A monocular advantage in responses to a particular category may reflect visual facilitation of responses for that category (*11*). Inputs from one eye are mostly segregated throughout the subcortical visual pathway (*27*), whereas inputs from the two eyes appear to be integrated to a greater extent in the extrastriate cortex (*28*). Two images presented to the same eye are likely to activate overlapping populations of monocular subcortical neurons, whereas two images presented to different eyes are not. In a previous study (*11*), the observation of monocular advantage for snakes and the absence of monocular advantage for guns indicate that visual facilitation of responses may be specific to evolutionary threats. Consistent with this previous study, we observed a monocular advantage for snakes but not for guns. Interestingly, the images of humanoid robots also showed a monocular advantage, potentially reflecting visual facilitation of responses for humanoid robots.

The amygdala, a subcortical structure in the anterior-temporal lobe, is located in an evolutionarily old part of the brain and is shared by other mammals. It is assumed to be the neural basis of hardwired “fear module” that allows us to automatically and quickly detect threatening stimuli (*10*). Studies have well documented that the amygdala responds selectively to evolutionary threats, irrespective of the affective valence, such as animate entities (*21, 29, 30*) and depictions of humans (*31-33*). Our results showing that no amygdala activity in response to conscious presentation of humanoid robot images seemingly support perception of humanoid robots as a modern threat. However, when automatic and controlled evaluations of threat sometimes differ, the more positive controlled processing can moderate more negative automatic processing (*15*). With the positive explicit attitude to humanoid robots found in present study, the absence of amygdala response to conscious presented humanoid robot images is understandable. Interestingly, although controlled processing can eliminate amygdala activity caused by consciously presented threats, the amygdala still shows greater responses to threats when threats were presented unconsciously (*15-17*). Our present study found that greater amygdala activity was induced by humanoid robot images compared to human images under unconscious presentation. These results potentially reflect the automaticity with rapid recruitment of amygdala for humanoid robot-related stimuli processing.

It has been proposed that there is enhanced resistance to extinction to evolutionary threats relative to modern threats (*10*). Modern threatening stimuli like pictures of guns have not induced the same resistance to extinction as do pictures of snakes (*19, 20*). Our study showed that associative learning successfully weakened participants’ negative implicit attitude to humanoid robots and showed that amygdala-hippocampus functional connectivity was significantly increased after this associative learning. Furthermore, the implicit attitude change was negatively correlated with the functional connectivity change in the extent of amygdala-hippocampus coupling. The amygdala processes emotional information and uses it for associative learning (*34*). The hippocampus is important for the consolidation of information, including short-term and long-term memory (*35*). The connectivity of amygdala-hippocampal circuit is understood to underlie the neural basis of emotion associative learning (*35, 36*). Our results indicate that a positive humanoid robot-related memory has been established, which is competitive with the priori negative humanoid robot-related memory. However, the amygdala could still automatically and quickly detect humanoid robots, indicating that the amygdala still regard humanoid robots as a threat.

An enormous amount of literature has focused on the psychological perspective for how humanoid robots are perceived. In many science fiction accounts, humanoid robots are presented as perfect soldiers who never tire or as ideal servants who always obey (*37*). In reality, people long balk at the idea that robots have any human-like mental capacities. But research emphasizes that people’s minds are matters of perception (*38*), and we perceive robots based on the ascriptions such as some ability to think, remember, and exert self-control (*39*). The more human-like a robot looks, the more people perceive it as having mental capacities, a phenomenon called anthropomorphism (*40*). Besides, the appearances of robots also convey meaning. For instance, people attribute species to some robots. Aibo is very clearly a robot dog, while Justo Cat is a robot cat. It is noteworthy that people possibly attribute race to humanoid robots (*41*). The appearances of these robots result that people perceive the robots as mechanical versions of these animals or humans. We are entering an age where there will likely be a new type of entity that combines some properties of machines with some apparently psychological capacities that were previously only evident in humans (*42*). Combined with our demonstration that humans may perceive humanoid robots as an evolutionary threat, future humanoid robots with superhuman intelligence may be perceived as a new “human race” who will fight for their own rights and enslave us. With humanoid robots, we should consider security and ethics as early as possible, and put these considerations into the technologies we develop.

In summary, this study demonstrates that humanoid robots are perceived as an evolutionary threat, even though humanoid robots are manmade, modern objects. Our findings provide new insights into the perception of humanoid robots from an evolutionary perspective, which can help to redefine human-robot interaction and inform robot ethics.

## Acknowledgements

This work was supported by grants from The National Key Basic Research Program (2018YFC0831101), The National Natural Science Foundation of China (71942003, 31771221, 61773360, and 71874170), Major Project of Philosophy and Social Science Research, Ministry of Education of China (19JZD010), CAS-VPST Silk Road Science Fund 2021 (GLHZ202128), Collaborative Innovation Program of Hefei Science Center, CAS (2020HSC-CIP001), and China Postdoctoral Science Foundation (2016M592051). A portion of the numerical calculations in this study were performed with the supercomputing system at the Supercomputing Centre of USTC.

## Author contributions

ZDW, YC, and XCZ conceived and designed the study. ZDW and YC obtained the findings. YC was responsible for acquisition of data. ZDW, YC analyzed and interpreted the data. PYZ, YP, JCR, RJZ, BSQ, YCB, SHH and DRZ provided administrative, technical, or material support. XCZ supervised the study. ZDW and YC drafted the paper and all authors contributed to critical revision for intellectual content.

## Data availability statement

The complete dataset is available from the corresponding author.

## Code availability statement

The MATLAB and AFNI code is available from the corresponding author.

