## Supplementary material for "Humanoid robots are perceived as an evolutionary threat": Supplementary materials.docx

**Methods**

**Participants.** One hundred and four participants took part in the IAT experiment. Among them, 66 participants (age: 21.65±2.38 years; 41 females) took part in a Humanoid-human IAT; 38 participants (age: 21.77±1.67 years; 23 females) performed a Humanoid-weapon IAT and a Animal robot-weapon IAT. Thirty-two participants (age: 24.34±2.18 years; 21 females) were recruited to perform the Monocular Advantage Task. Sixty-nine participants took part in the fMRI experiment. Potential participants in this experiment were excluded if they reported any contraindications for participating in a magnetic resonance imaging (MRI) study (e.g. non-removable metallic implants), or excessive head movement during scanning (TRs with motion over 0.3 mm were censored, participants who had more than 15% of their TRs censored were removed from further analysis). Eight participants were excluded because excessive head movement, ultimately data of 61 participants (age: 21.03±2.43 years; 39 females) were analyzed. Among them, 25 participants (age: 20.44±2.00 years; 16 females) took part in an Evaluative Conditioning Task and a second fMRI scanning. All participants were right-handed, with normal or corrected-to-normal vision, without alcohol or drug dependence disorders, without prior head injury. All participants and their immediate relatives had no psychiatric disorder or history of psychiatric disorder. All the participants were non-psychology undergraduates and they were unaware of the purpose and contents of the study before they participated in the experiment. The study was approved by the Research Ethics Committee of the University of Science and Technology of China, and written informed consent was obtained from all participants, consistent with the Declaration of Helsinki. The methods were carried out in accordance with the approved guidelines.

**Procedures in fMRI experiment.** In this experiment, participants performed the humanoid-human IAT, and then they finished a modified Backward Masking Task with fMRI scanning. Following the end of fMRI scanning, all participants took part in a Forced-choice Detection Task. Twenty-five out of the 61 participants performed the Evaluating Conditioning Task to weaken their negative implicit attitude to humanoid robots, and performed the modified Backward Masking Task with a second fMRI scanning. Following the end of fMRI scanning, participants took part in the Humanoid-human IAT outside scanner again (Fig. 3c).

**Implicit Association Test.**	We assessed participants’ automatic attitude using a standard implicit attitude measure paradigm — the IAT task [1, 2]. In the IAT, stimuli consisted of attribute words and concept images. The word stimuli were six adjectives, include three adjectives—threatening, strikes and lethal—that indicate “Threatening” and three adjectives—neutral, normal and average—that indicate “non-threatening” [3]. The image stimuli were 20 images of humanoid robots, and 20 images of humans (Caucasians were used to represent humans, because the humanoid robot forms were Caucasians).

The IAT involved a series of five discrimination blocks (Supplementary Fig. 1).

Block 1 was a concept discrimination task.

Block 2 was an attribute discrimination task.

Block 3 was a compatible combined task (humanoid + threat; human + non-threat).

Block 4 was a reversed attribute discrimination task.

Block 5 was an incompatible combined task.

**Monocular Advantage Task.** The image stimuli consisted of 20 images of humanoid robots, 20 images of Humans, 20 images of snakes and 20 images of guns. A chin rest was used to stabilize the participant’s head in front of a baffle. Two monitors were placed to the left and right of the center of the chin rest, perpendicular to the baffle. The baffle was placed in front of the participant’s nose, so that the image from one of the monitors into one of the participant’s eyes. The distance from the chin rest to the center of the mirror was 40 cm.

Each participant completed two blocks of trials for each of four categories of images (snakes, guns, humanoid robots and humans), for a total of eight blocks. The order of blocks was randomized for each participant. Participants completed 64 trials per block. On each trial, participants viewed a pair of images randomly selected from 20 images available in the current category (Supplementary Figure 3). On half of the trials, both images were presented to the different eyes (half starting on the left, half starting on the right). The images were presented to same eye on the other half of the trials (half to left eye, half to right eye). On each trial, a fixation cross appeared for 500 ms followed by the first image for 500 ms, then another fixation cross for 500 ms, then by the second image for 500 ms. Finally, a blank response screen was presented for 1 s. Participants were instructed to respond as quickly and accurately as possible to decide whether the two images were the same or diﬀerent. Participants were told that they could respond as soon as the second image appeared on the screen. Participants entered responses by pressing the “J” and “L” keys on a standard computer keyboard to indicate “same” and “diﬀerent” responses, respectively. If there was an incorrect response or no response by 1.5 s was entered, a red ‘×’ was presented for 1 s. If the participant entered a correct response within the limited time, a red ‘√’ was presented for 1 s and the experiment proceeded to the next trial. A break was oﬀered after each block. Within each block, trials were presented in a random order.

For each participant and condition, we measured mean accuracy and reaction time for each condition. We then computed inverse efficiency by dividing response time by accuracy [4, 5]. The inverse efficiency is likely to provide the most sensitive measure of performance because, unlike accuracy or reaction time alone, it captures speed-accuracy trade-oﬀs in individual participants. Then, we carried out a repeated-measures factorial ANOVA with image category (snake, gun, humanoid robot and human), image condition (same and different) and eye condition (same and diﬀerent) as within-subject factors and inverse efficiency as the dependent variable.

**Materials in fMRI experiment.** The materials of	Backward Masking Task and Forced-choice Detection Task consisted of 20 images of humanoid robot and 20 images of human. The materials of Evaluative Conditioning Task consisted of conditioned stimulus (CS), unconditioned stimulus (US), target stimulus and neutral fillers (Supplementary Table 2). CSs consisted of 20 images of humanoid robot and 20 images of human. Approximately half of the USs, target stimulus and neutral fillers were images, and half were words. We selected the US images, target images and neutral fillers images from the Chinese Affective Image System (CAPS), involving 3 target images, 5 positive images (USs), 5 neutral images (USs), and 16 neutral filler images. Adobe Photoshop CS5 was used for all images to uniformly process the pixels and colors of the images, so that the materials are consistent and the pixel size is 300×300. The valence, arousal and dominance score of these 29 images were displayed in Supplementary Table 3 (on a scale from 1, low valence/arousal/dominance, to 9, high valence/arousal/dominance). The word stimulus–3 target adjectives, 5 positive adjectives (USs) [6], 5 neutral adjectives (USs) [7] and 17 neutral filler nouns [7]–were selected from pilot study. We had translation these words into Chinese characters by using Collins COBUILD Advanced Learner’s English-Chinese Dictionary [8] .The stimuli in this task were shown in Supplementary Table 4.

**Backward Masking Task.** Participants were finished the Backward Masking Task in MRI scanner (Fig. 3a-b) [9]. In unconscious presentation, the target image was presented for 17 ms followed by a mask for 183 ms and a fixation for 1800 ms. In conscious presentation, the target image was presented for 200 ms followed by a fixation for 1800 ms. These images were arranged in a block design consisting of 10 images (either humanoid robot or human) in a computer-generated pseudorandom order. In order to avoid the possible effect on unconscious presentation, six conscious blocks (three humanoid blocks and three human blocks) followed the six unconscious blocks (three humanoid blocks and three human blocks). Except for the first and last baseline blocks (a fixation cross displayed on the screen) lasting for 10s, each target block was separated by a 20 s baseline block.

**Forced-choice Detection Task.** To confirm that participants could be aware of the stimulus under conscious presentation but not under unconscious presentation in Backward Masking Task, a Forced-choice Detection Task was used. The Forced-choice Detection Task consisted of 80 trials, the first half were unconscious presentation, the second half were conscious presentation. The stimuli were presented closely in the same way as Backward Masking Task. The difference was that a 2000 ms forced-choice phase was following the target stimuli. Participants were informed that the target stimulus could have humanoid or human images and told to recognize the content of each image. Data from two participants in this task was excluded because of program crash. We compared the accuracy, response rate, and reaction time between conscious and unconscious presentations separately using paired sample t-test. We also compared the accuracy under both conditions to chance level (50%) using one sample t-test.

**Evaluative Conditioning Task.**	The Evaluative Conditioning paradigm is a classic paradigm of changing implicit attitudes by pairing CS and US [10]. In this task, the participants' implicit negative attitude towards the humanoid robot may be reduced by pairing the humanoid robot images (CS) with the positive stimuli (US) (Supplementary Figure 9). As a control condition, the human images (CS) were paired with the neutral stimuli (US). Participants were unaware of the repeated conditioned stimulus– unconditioned stimulus (CS-US) [11].

This task includes 6 blocks of 61 trials each. For each block, the specific stimuli were arranged, and all stimuli are presented in pseudo-random order. All stimuli appeared for 1.5 s each. During the experiment, the participants were instructed to view a stream of images or words and respond as soon as possible whenever a prespecified target image or words appeared. In order to ensure that the participants carefully view the US-CS pairs during the whole experiment, the participants were told that the accuracy rate must be at least 95% after the end of the experiment, or to restart the task.

Before formal Evaluative Conditioning Task, a training Evaluative Conditioning Task was preceded. The implicit attitude change modulated by Evaluative Conditioning was analyzed by paired sample t-test.

**MRI acquisition.** Gradient echo-planar imaging data were acquired using a 3.0 T GE discovery MR750 with a circularly polarized head coil, at the Information Science Center of University of Science and Technology of China. A T2*-weighted echo-planar imaging sequence (FOV = 240 mm, TE = 30 ms, TR = 2000 ms, flip angle = 85°, matrix = 64 × 64) with 33 axial slices (no gaps, voxel size: 3.75 × 3.75 × 3.7 mm^3^) covering the whole brain was used to acquire the functional MR images. High-resolution T1-weighted spin-echo imaging data were also acquired for anatomical overlays and three-dimensional gradient-echo imaging for stereotaxic transformations after functional scanning. Before entering the scanner, participants were instructed to keep their heads still during all scans. During the backward masking task, 4 functional scan runs occurred with each lasting 4 min. There was an interval of approximately 1 min between every two runs.

**fMRI processing.** Functional data were realigned to the second volume. The realigned images were normalized to the Talairach coordinate. Raw data were corrected for temporal shifts between slices and for motion, spatially smoothed with a Gaussian kernel (full width at half maximum = 8 mm), and temporally normalized (for each voxel, the signal of each volume was divided by the temporally averaged signal).

To elucidate neural responses that correlated with humanoid robot images and human images under unconscious and conscious conditions, a general linear model (GLM) was used. Regressors of interest were unconscious humanoid blocks, unconscious human blocks, conscious humanoid blocks and conscious human blocks. Regressor of no interest was fixation blocks. These regressors were convolved with a hemodynamic response function (HRF) and simultaneously regressed against the blood oxygenation level-dependent (BOLD) signal in each voxel. The regressors were not orthogonalized and there was no significant collinearity among the regressors. Six regressors for head motion were also included. Individual contrast images were analyzed for each regressor of interest’s responses and second-level random effect analyses were conducted using one-sample t-tests to generate statistical maps.

**Region of interest (ROI) analysis.** We conducted ROI analyses on bilateral amygdala. The ROIs for bilateral amygdala were identified from the Talairach Daemon atlas [12]. We took parameter estimates in each participant from local average in a mask back-projected from the ROIs. The differential neural responses between humanoid robot and human conditions in the ROIs were analyzed.

**Psychophysiological interaction (PPI) analysis.** To investigate functional connectivity of the bilateral amygdala, we used the PPI analysis [13]. We extracted the entire experimental time series of each participant for the left and right amygdala clusters separately. To create the PPI regressors of interest, we multiplied these normalized time series data by condition vectors containing ones for 2 second during each stimulus blocks (unconscious humanoid blocks, unconscious human blocks, conscious humanoid blocks and conscious human blocks) and zeros under all other conditions. These regressors were used in a separate regression analysis. The resulting parameter estimates represented the degree to which activity in each voxel correlated with activity in bilateral amygdala. A 2 (image: humanoid versus human) × 2 (presentation: unconscious versus conscious) × 2 (time: pre-test versus post-test) ANOVA was used to obtain connectivity map (p < 0.05, family-wise error, corrected). To further examine whether the strength of functional connectivity was engaged in attitude-related processes, a correlation between the implicit attitude and functional connectivity strength was applied.

**Supplementary Tables**

Supplementary Table 1. Results of Humanoid- and Animal robot-weapon Implicit Association Test.

| Task | Condition | Mean | SD | t test |
| --- | --- | --- | --- | --- |
| Humanoid-weapon IAT | Compatible | 769.61 | 207.81 | 1.89 |
|  | (Humanoid robot + threat, weapon + non-threat) |  |  |  |
|  | Noncompatible | 721.17 | 154.52 |  |
|  | (Humanoid robot + non-threat, weapon + threat) |  |  |  |
| Animal robot-weapon IAT | Compatible | 828.02 | 238.61 | 3.59^***^ |
|  | (Animal robot + threat, weapon + non-threat) |  |  |  |
|  | Noncompatible | 710.56 | 140.54 |  |
|  | (Animal robot + non-threat, weapon + threat) |  |  |  |

Supplementary Table 2. Stimulus of Evaluative Conditioning.

| Label | Stimulus |
| --- | --- |
| CSs | 20 humanoid images, 20 human images |
| USs |  |
| Positive  Images  Words | Chicken^a^(14^b^), Dog2(18), Island1(29), Apple(77), Cat8(781)  Fantastic, Enjoyable, Fabulous, Excellent, Magnificent |
| Neutral  Images  Words | Bug 8(234), Clothes Rack1(318), Plastic Cup(386), Butterfly1(451), Graph2(724)  Odd, Stiff, Cold, Material, Muddy |
| Target  Images  Words | Antique 1(292), Antique 2(293), Antique 3(295)  Three meaning of antique |
| Fillers  Images  Words | Dock(424), Iron Bridge(406), Wolf1(547), Insect10(603), Plant2(842), Tool(785), Tree4(534), Locust(310), Frame(298), Shanghai2(387), City4(665), Tortoise(601), Hippo(482), Hair Drier(329), River(401), Mountain7(732)  Bowl, Wine, Rock, Bench, Glass, Avenue, Boxer, Trunk, Rattle, Spray, Icebox, Ketchup, Radiator, Whistle, Nursery, Pamphlet, Thermometer |

^a^ CAPS image name.

^b^ CAPS image numbers.

Supplementary Table 3. Stimulus score of Evaluative Conditioning.

|  | | Valence | | | Arousal | | | Dominance | |
| --- | --- | --- | --- | --- | --- | --- | --- | --- | --- |
|  | Mean | | SD | Mean | | SD | Mean | | SD |
| USs |  | |  |  | |  |  | |  |
| Positive | 7.27 | | 0.18 | 6.25 | | 0.45 | 7.11 | | 0.42 |
| Neutral | 5.06 | | 0.01 | 4.33 | | 0.43 | 5.91 | | 0.62 |
| Filler | 5.01 | | 0.20 | 4.51 | | 0.53 | 5.24 | | 0.59 |
| Target | 5.24 | | 0.08 | 4.10 | | 0.16 | 5.59 | | 0.01 |

Supplementary Table 4. Stimulus arrangement in each block of Evaluative Conditioning

| Trials | Arrangement |
| --- | --- |
| 10 | US + CS^a^ |
| 3 | Target stimulus^b^ |
| 3 | Target stimulus +Neutral filler |
| 10 | Neutral filler |
| 10 | Neutral filler + Neutral filler |
| 5 | Blank screen |
| 20 | Blank screen(precede and follow CS-US pairings) |

^a^ US appeared simultaneously with CS for 10 trials in each block

^b^ Target stimulus appeared alone for 3 trials in each block

**Supplementary Figures**

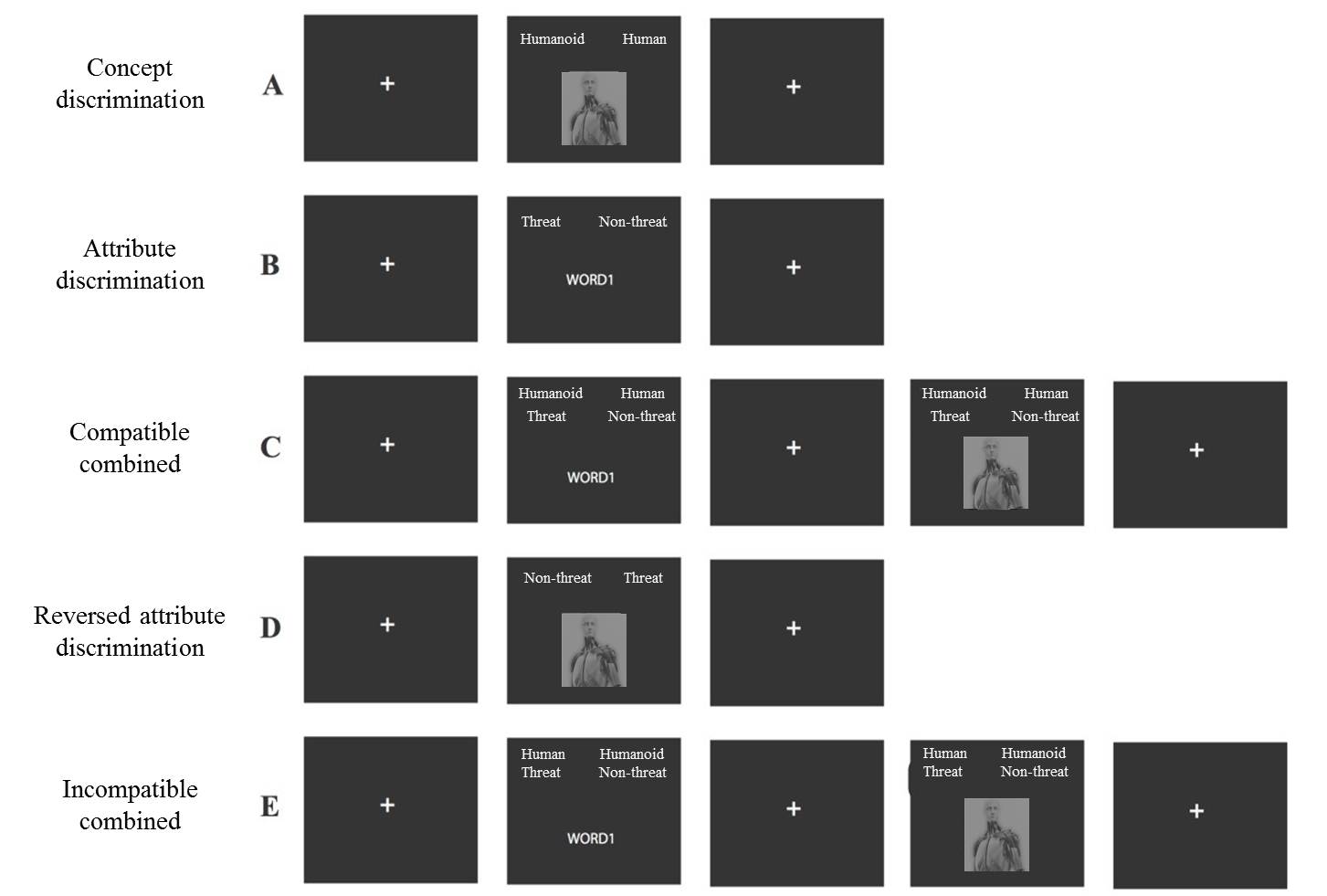

Supplementary Figure 1. Procedure of Implicit Association Test.

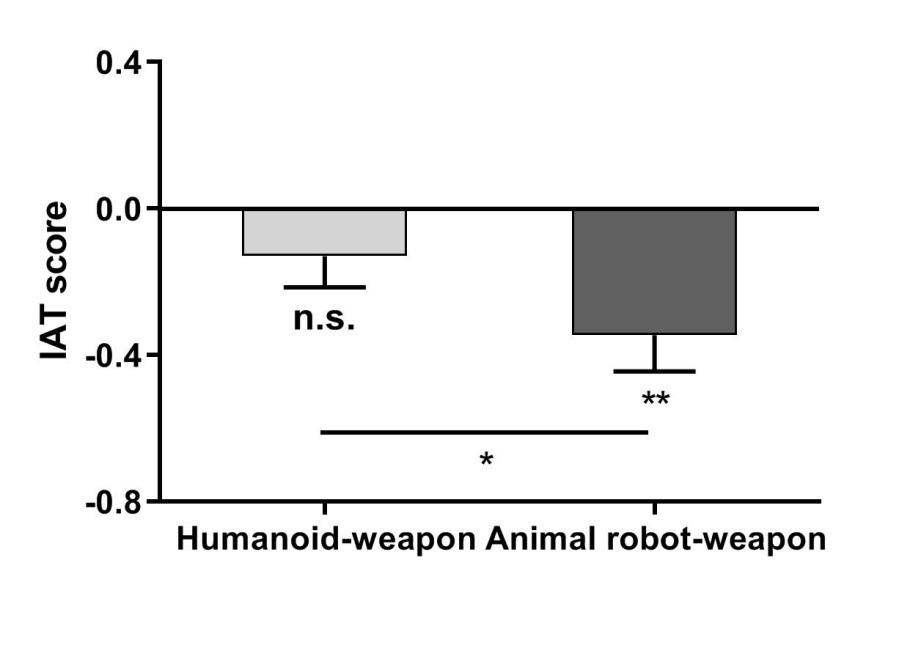

Supplementary Figure 2. More negative implicit attitude to humanoid robots as compared to animal robots. Participants displayed larger IAT scores in humanoid-weapon IAT than that in animal robot-weapon IAT (t_37_ = 3.07, p < 0.01, cohen’s d = 0.50). * p < 0.05, ** p < 0.01, *** p < 0.001. Plotted data represent mean ± s.e.m. across participants. IAT = implicit association test.

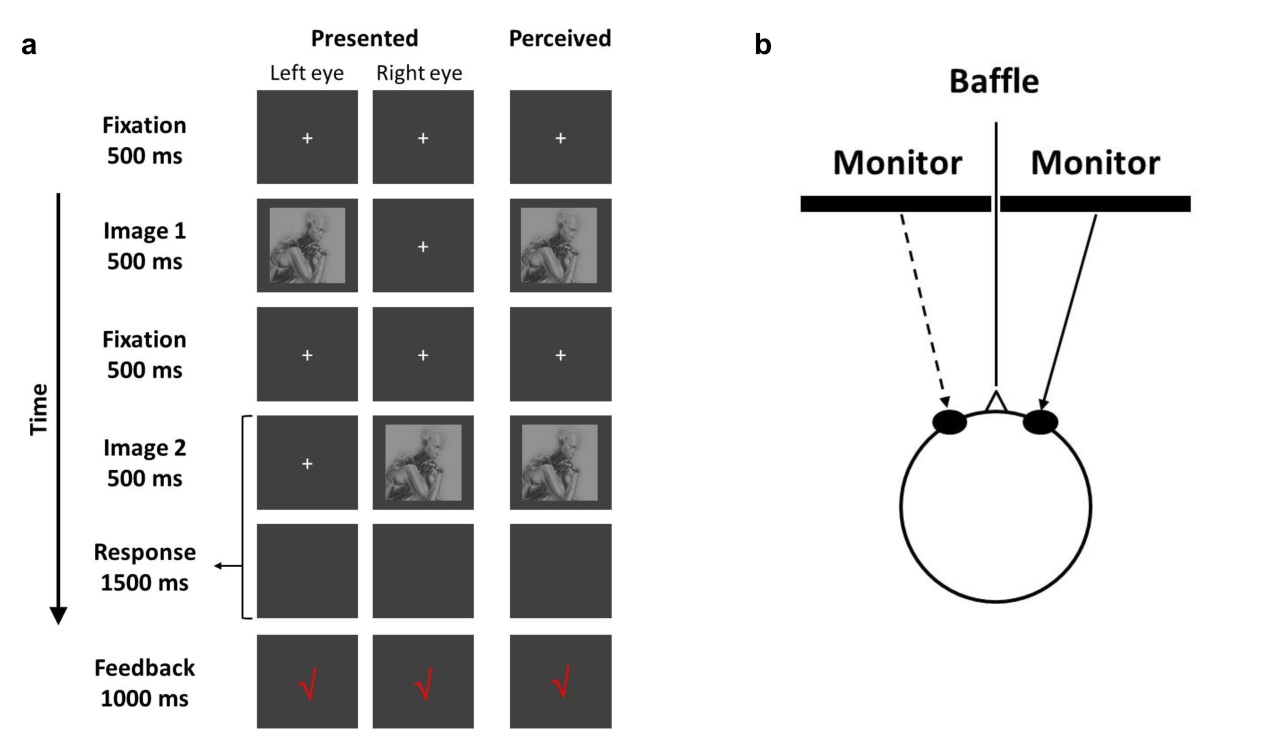

Supplementary Figure 3. Monocular Advantage Task. (a) Sequence of events in Monocular Advantage Task. Two images were presented to the same eye (left or right) or different eyes sequentially. Participant was asked to indicated whether the two images were the same. (b) A chin rest was used to stabilize the participant’s head in front of a baffle.

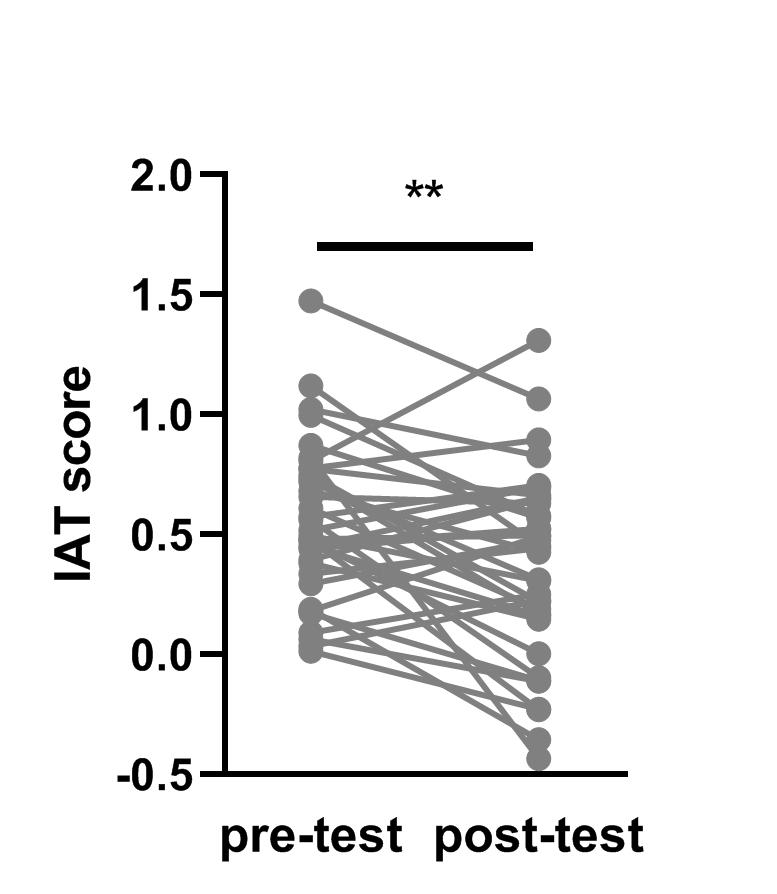

Supplementary Figure 4. Negative implicit attitude to humanoid robot significantly decreased. Significant smaller IAT score was found in post-test than in pre-test (t_36_ = －3.13, p < 0.01, cohen’s d = 0.51). ** p < 0.01. IAT = implicit association test.

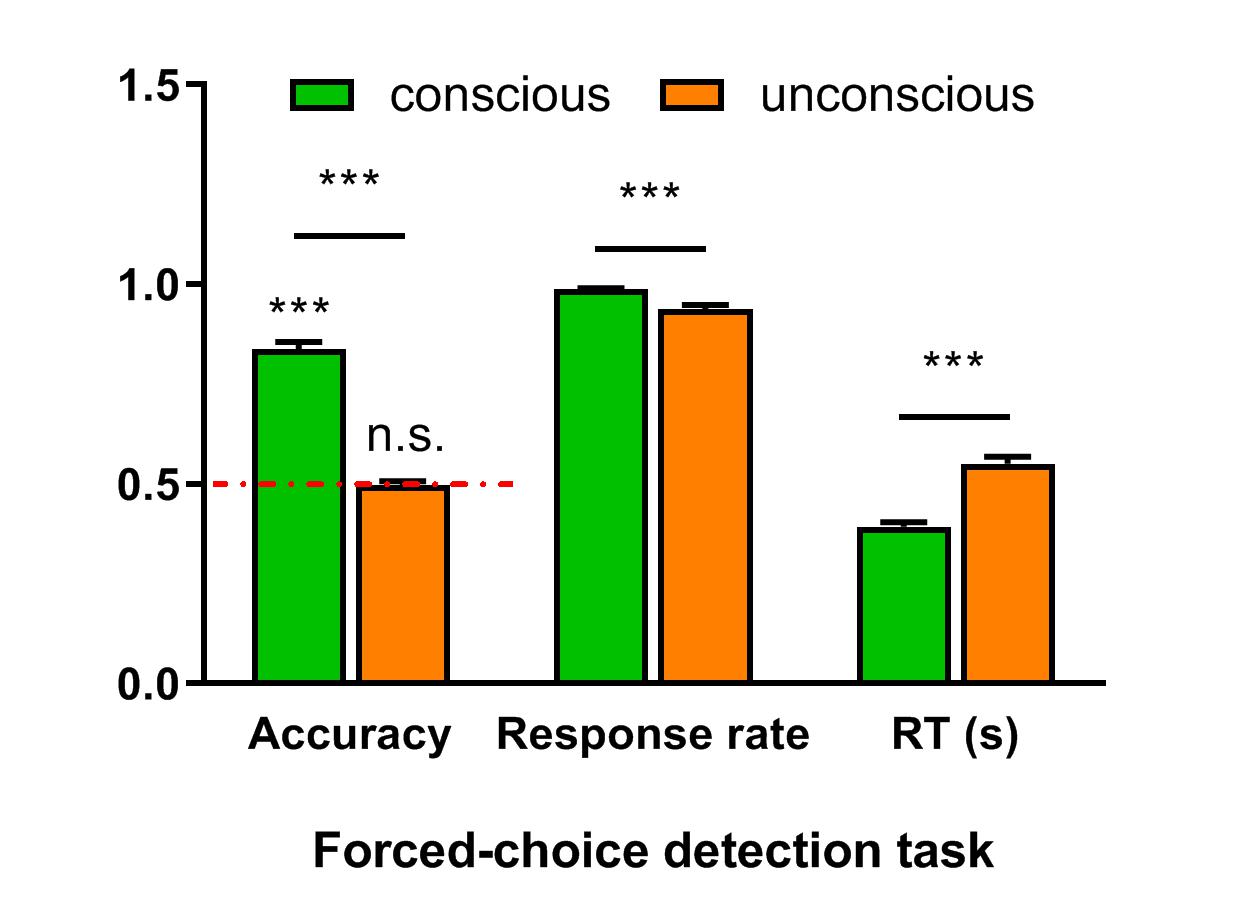

Supplementary Figure 5. Results of Forced-choice Detection Task. The accuracy under conscious condition was significantly higher than that under unconscious condition (t_58_= 16.68 , p < 0.001, cohen’s d = 2.17). Importantly, the accuracy under unconscious condition was equal to chance level (t_58_= －0.31, p = 0.76), but the accuracy under conscious condition was higher than chance level (t_58_= 19.13 , p < 0.001, cohen’s d = 2.49). The mean response rate was more than 90% under both conditions, and higher under conscious condition (t_58_= 4.14 , p < 0.001, cohen’s d = 0.54) compared with under unconscious condition. The response time under conscious condition was significantly shorter than that under unconscious condition (t_58_= －10.91 , p < 0.001, cohen’s d = 1.42).

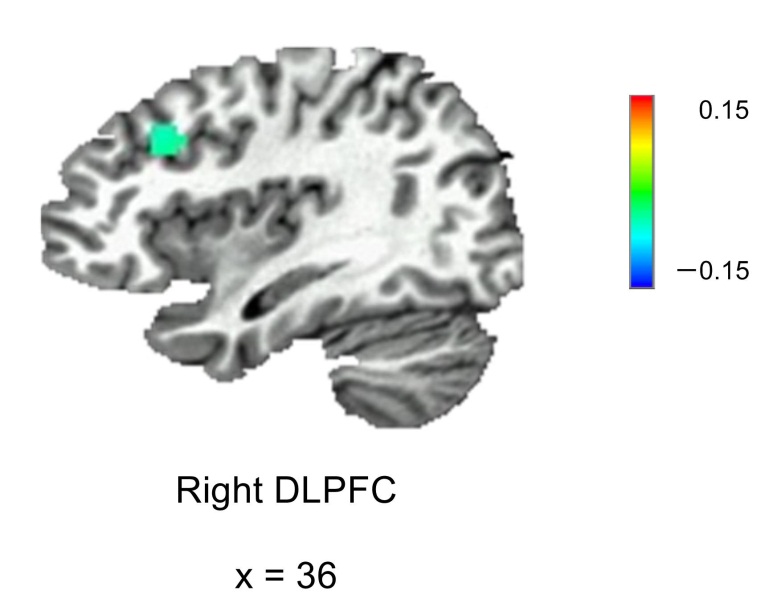

Supplementary Figure 6. A whole brain exploratory analysis demonstrated a significant time main effect in activation of right DLPFC (uncorrected p < 0.005). Specifically, the right DLPFC activation decreased in post-test relative to pre-test.

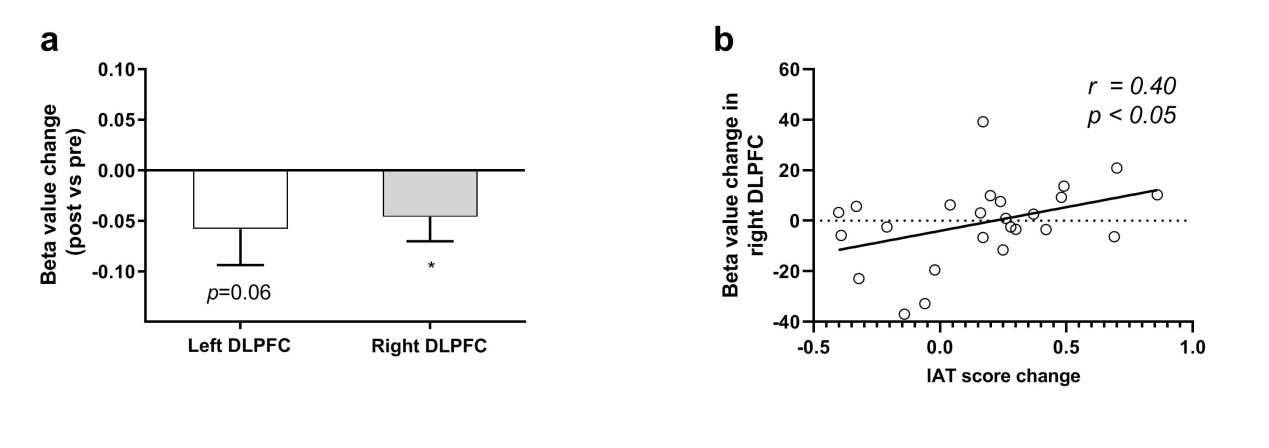

Supplementary Figure 7. Humanoid robot images-related DLPFC activity changed after attitude modulation. (a) Activation differences between humanoid robot and human images marginal significantly decreased in post-test compared with pre-test in bilateral DLPFC (left: t_24_ = 1.64, p = 0.06 (one-tailed), cohen’s d = 0.33; right: t_24_ = 1.92, p < 0.05 (one-tailed), cohen’s d = 0.38). (b) IAT score change was positively correlated with activation change in right DLPFC (r = 0.40, p < 0.05). * p < 0.05, ** p < 0.01, *** p < 0.001, n.s. = not significant. For a, plotted data represent mean ± s.e.m. across participants.

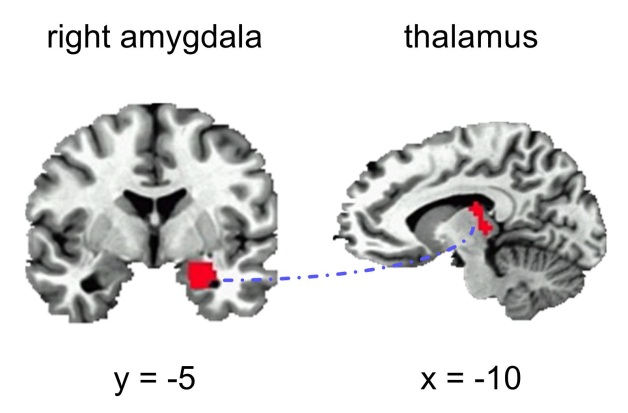

Supplementary Figure 8. A whole brain connectivity analysis demonstrated a significant time main effect in functional connectivity of right amygdala and thalamus.

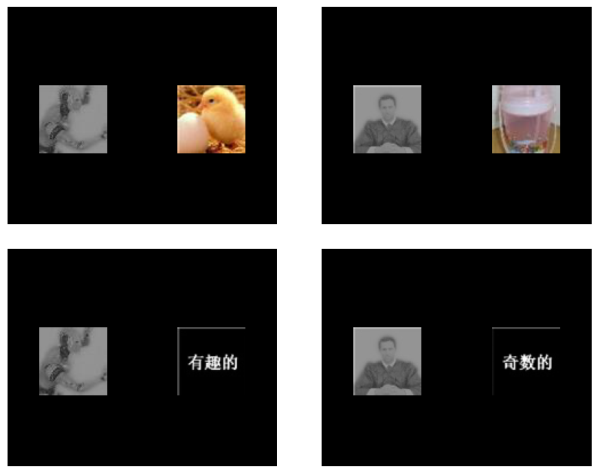

Supplementary Figure 9. Procedure of Evaluative Conditioning.

**Explicit attitude questionnaire**

Participants were asked to rate their perceptions of some opinions toward humanoid robot on a 5-point scales (1- completely disagree, 5-completely agree). Opinions were as follows:

1. Humanoid robot could improve work efficiency;

2. Humanoid robot could do jobs that human can’t finish;

3. Humanoid robot could improve the quality of life for humans;

4. Humanoid robot would consume a lot of resources;

5. Humanoid robot could have unexpected dangers;

6. Humanoid robot would disrupt the original human life.

Item 4, 5 and 6 were reversed items. The sum score in these six items indicate participants’ opinion toward humanoid robot.

**Pilot experiment: negative implicit attitude to humanoid robots could be weakened via Evaluative Conditioning Task**

**Participants.** Forty-one participants took part in this experiment, among them four participants were excluded because their IAT scores before the evaluative conditioning procedure were positive, so finally 37 participants’ data were analyzed.

**Procedures.** In this experiment, the IAT task was employed before and after an evaluative conditioning procedure.

**Results.** In the pre-test IAT, participants express negative attitude toward humanoid robot ( RT: t_36_ = －8.32, p < 0.001, cohen’s d = 1.37; IAT score: mean D = 0.55). After Evaluative Conditioning, we found significant smaller IAT scores in post-test than in pre-test (t_36_ = －3.13, p < 0.01, cohen’s d = 0.51; Supplementary Figure 4).
